# Novel PE and APC Tandems: Additional Near-Infrared Fluorochromes for Use in Spectral Flow Cytometry

**DOI:** 10.1101/2020.06.25.165381

**Authors:** Yekyung Seong, Yian Wu, Denny Nguyen, Archana Thakur, Tuan Andrew Nguyen, Fiona Harding

**Author notes:** Corresponding author, 1500 Seaport Blvd, Redwood City, CA 94063.

## Abstract

Recent advances in flow cytometry instrumentation and fluorochrome chemistries have greatly increased fluorescent conjugated antibody combinations that can be used reliably and easily in routine experiments. The Cytek Aurora flow cytometer has three excitation lasers (405nm, 488nm, and 640nm) and incorporates the latest Avalanche Photodiode (APD) technology, demonstrating significant improvement in sensitivity for fluorescent emission signals longer than 800nm. However, there are limited commercially available fluorochromes capable of excitation with peak emission signals beyond 800nm. To address this gap, we engineered 6 new fluorochromes: PE-750, PE-800, PE-830 for the 488nm laser and APC-750, APC-800, APC-830 for the 640nm laser. Utilizing the principal of fluorescence resonance energy transfer (FRET), these novel structures were created by covalently linking a protein donor dye with an organic small molecule acceptor dye. Additionally, each of these fluorochrome conjugates were shown to be compatible with Fixation/Permeabilization buffer reagents, and demonstrated acceptable brightness and stability when conjugated to antigen-specific monoclonal antibodies. These six novel fluorochrome reagents can increase the numbers of fluorochromes that can be used on a 3-laser spectral flow cytometer.

## INTRODUCTION

An advantage of near-infrared emission (wavelengths from 780nm to 900nm) is the limited interference from cellular autofluorescence sometimes associated with shorter wavelengths (400nm to 550nm)^1^. As a result, the use of fluorochromes with near-infrared emissions may result in higher sensitivity and improved fluorescent staining indexes. Near-infrared emission fluorochromes also tend to introduce minimal spillover into detectors designated for fluorochromes with shorter wavelength emission spectra.

Despite these advantages, there has been limited application of fluorochromes with near-infrared emission spectra in flow cytometry. Most commercial cytometers are commonly manufactured with a 488nm blue laser and 640nm red laser, but not a near-infrared laser. To make fluorochromes with peak excitation in the 488nm to 640nm range and peak emission within the near-infrared region, manufacturers rely on the principle of FRET, coupling a donor base fluorochrome (e.g. PE or APC) with an acceptor fluorochrome that emits near-infrared fluorescence (e.g. Cy7)^2^. These are commonly referred to as “tandem dyes”. Historically, PE-Cy7 and APC-Cy7 are among the most widely used tandem dyes with peak emission wavelengths near 780nm. Up until recently, there were no other commercial PE and APC tandem fluorochromes with emission wavelengths longer than 780nm that could be easily distinguished from PE-Cy7 and APC-Cy7. Very low quantum efficiency of photomultiplier tube (PMT) detector for emissions longer than 800nm could explain the limited commercial availability of fluorochromes with near-infrared emission in flow cytometry^3^. Recent advances in optic technology led to the commercial adoption of Avalanche Photodiodes (APD) (e.g., Beckman Coulter Cytoflex and Cytek Aurora), which have improved quantum efficiencies over photomultiplier tubes for fluorescent emission in the infrared region^4^.

In this technical note, we describe and characterize the performance of the following six novel fluorochromes: PE-750, PE-800, PE-830, APC-750, APC-800, and APC-830 with peak emission wavelengths at 750nm, 800nm, 830nm, respectively.

## MATERIALS AND METHODS

### Subjects and Samples

The study was approved by the Institutional Review Board at Abbvie Biotherapeutics. The human blood samples were collected from healthy donors who registered for AbbVie Biotherapeutics Employee Blood Collection Program in Redwood City, CA. Whole blood was obtained in heparin-anticoagulated tubes (BD Biosciences, San José, CA) and then processed for staining on the same day of collection.

### Preparation of PE- and APC-linked fluorochromes

The conjugation method was adapted from the previously described protocol^5^. Prior to conjugation, R-Phycoerythrin (R-PE, 240kDa) (Prozyme, Hayward, CA) was extensively dialyzed into phosphate-buffered saline (PBS, pH 7.2) (GE Life Sciences, PA) using the Slide-A-Lyzer Dialysis Cassettes (Thermo Fisher, Carlsbad, CA) according to the manufacturer’s instructions. Briefly, the R-PE is dialyzed for 2 hours at room temperature and this process is repeated with fresh PBS for another 2 hours. The dialysis buffer was then replaced with fresh PBS, and the dialysis continued overnight at 4°C. The final concentration of dialyzed R-PE was 5.54 - 12.2 mg/ml. Lyophilized cross-linked Allophycocyanin (APC, 105kDa) (AAT Bioquest, Sunnyvale, CA) was resuspended with PBS to a final concentration of 2.5 mg/ml.

Organic small molecule NHS Ester fluorophores such as Dy704-P4, Dy704 (Dyomics, Germany), Dy800-P4 (Thermo Fisher, Carlsbad, CA), and iFluor810 (AAT Bioquest, Sunnyvale, CA) were dissolved with anhydrous DMSO (Thermo Fisher, Carlsbad, CA) to a final concentration of 1,000 nmol/ml. For example, 1mg of Dy800-P4 NHS Ester was dissolved in 952μl of anhydrous DMSO. Meanwhile, 1M sodium bicarbonate (pH 8.3-8.5) was prepared by dissolving sodium bicarbonate (Sigma-Aldrich, St Louis, MO) in deionized water.

The reaction condition is summarized in Table 1. The reaction was rotated at room temperature for at least 60 minutes. The absorbance spectra of the fluorochromes were preliminarily measured with SpectraMax M5 microplate reader (VWR, Radnor, PA) using 1:100 dilution in PBS. Desalting procedure is followed using the Zeba Spin 7K MWCO desalting columns (Thermo Fisher, Carlsbad, CA) according to the manufacturer’s instruction.

**Table 1.**
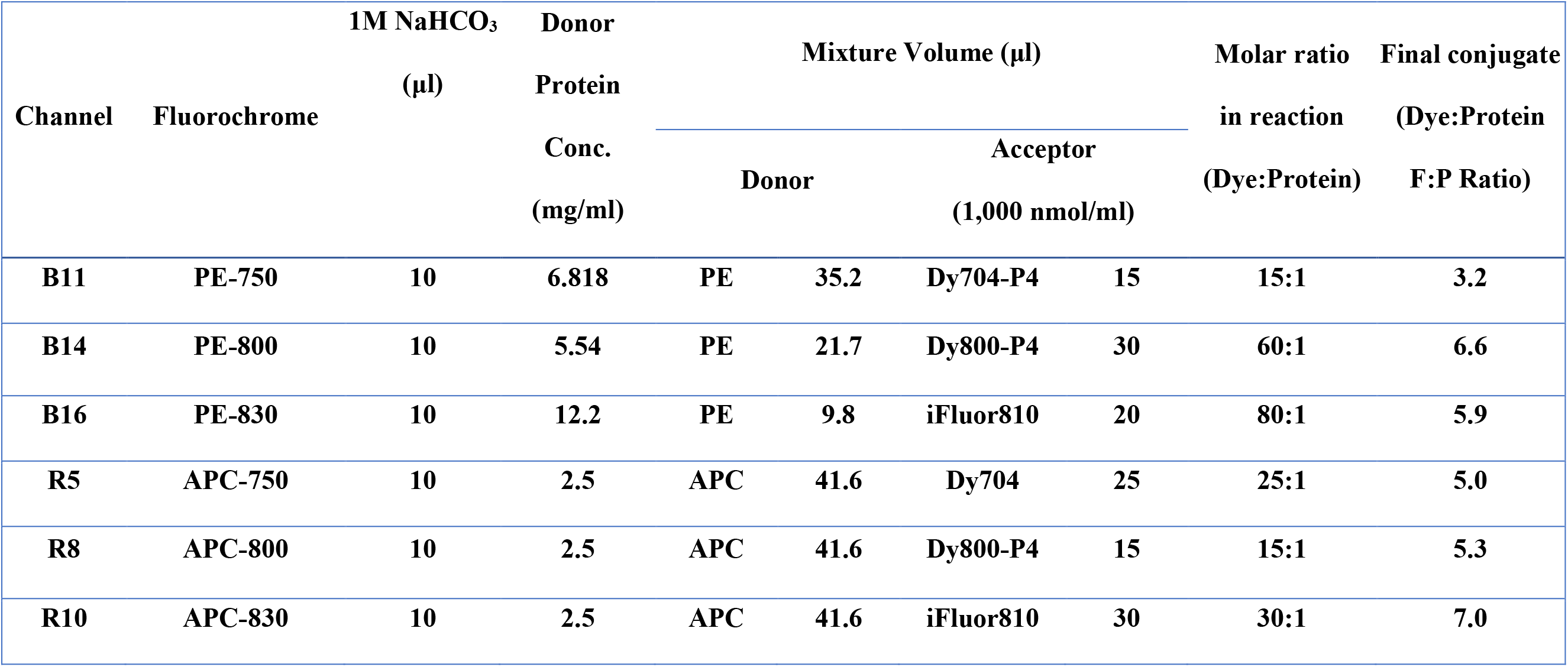
Reaction condition for pairs of a donor protein and an acceptor small molecule dye.

### Characterize the brightness of the tandem fluorochrome molecules

Anti-mouse Ig, κ compensation particles (BD Biosciences, San José, CA) were stained with 1μg of purified anti-Phycoerythrin antibody (clone PE001, BioLegend, La Jolla, CA), and then washed twice with staining buffer (1X PBS with 0.5% BSA). The compensation particles were then stained with PE tandem fluorochrome molecules.

Similarly, anti-mouse Ig, κ compensation particles were stained with 1μg of purified anti-Allophycocyanin antibody (clone APC003, BioLegend), and then washed twice with staining buffer. The compensation particles were then stained with APC tandem fluorochrome molecules. Stained samples were acquired on the spectral flow cytometer Aurora (Cytek, Fremont, CA).

### Click-chemistry to generate novel fluorochromes-conjugated antibodies

Conjugation of a monoclonal antibody and the tandem fluorochrome was achieved by click chemistry reaction between methyl-tetrazine and trans-cyclooctene-tetrazine (TCO) (Click Chemistry, Scottsdale, AZ). Click Chemistry works well in conjugation with tandem dyes because the conjugation can occur at near neutral pH conditions (pH = 7.2). TCO was tagged to 100 μg of each antibody. Each of the anti-human CD3 (Clone UCHT1), CD8α (Clone OKT-8), CD19 (Clone 4G7), CD20 (Clone 2H7) (Bio X Cell, West Lebanon, NH), CD16 (Clone 3G8, Leinco Technologies, Fenton, MO), and CD45 (Clone HI30, BioLegend) antibodies were mixed with 5 μl (5% of the total reaction volume) of 1M NaHCO_3_ with 100 μl of the PBS-based solution. Then, 20 nmol of TCO-PEG4-NHS ester was added to the mixture.

In the same manner, methyl-tetrazine was tagged to the fluorochromes. 100 μg of tandem fluorochromes were mixed with 5 μl of 1M NaHCO_3_ with 100 μl of the PBS-based solution. Then, 20 nmol of methyl-tetrazine-PEG4-NHS ester was added to the mixture. Those reaction mixtures were kept at room temperature for 60 minutes. Desalting procedure is followed for both mixtures using spin desalting columns (Thermo Fisher, Carlsbad, CA). The recovery protein amount after desalting was calculated as ~75 μg.

Cross-linking reaction was initiated by mixing the two reaction mixtures. Antibody-TCO was mixed with fluorochrome-methyl-tetrazine ester in 1:2 molar ratio. The reaction mixture was stored at 4°C overnight. The next day, protein stabilizing cocktails (Thermo Fisher, Carlsbad, CA) and bovine serum albumin were added. The final products were stored at −20°C.

Large-scale conjugation was done by AAT Bioquest for PE-750, APC-750 and by BioLegend for PE-800, PE-830, APC-800, and APC-830.

### Conjugation of antibodies to commercially available fluorochromes

Prior to the conjugation procedure, concentration of the antibodies should be higher than 1 mg/ml for an optimal reaction condition. If needed, monoclonal antibodies were concentrated by the Amicon Ultra centrifugal filter (Millipore, Burlington, MA).

Anti-human CCR6 (Clone G034E3, BioLegend) was conjugated with CF680 (Biotium, Fremont, CA) in 1:10 molar ratio. 1M sodium bicarbonate (pH 8.3-8.5) was added to 10% of total volume. After 2-hour incubation at room temperature, the mixture went through the desalting spin column.

Anti-human CCR3 (Clone 5E8, BioLegend) was conjugated to biotin using EZ-Link Sulfo-NHS-LC-Biotin (Thermo Fisher, Carlsbad, CA). 2 mg of antibody was mixed with 150 μg of biotin reagent. The mixture was incubated at room temperature for 30 minutes and subsequently desalted.

Anti-human CD11b (Clone M1/70, Leinco Technologies, Fenton, MO) was conjugated with Alexa Fluor 532 NHS ester according to the manufacturer’s instructions. 0.75 nmol of CD11b antibody was added to the pre-mix and 1M sodium bicarbonate (pH 8.3-8.5) was added to 10% of total volume. After 2-hour incubation at room temperature, the mixture went through the desalting spin column.

### Comparison of sensitivity between PMT-based and APD-based flow cytometers

PMT-based flow cytometer was compared against APD-based flow cytometer to evaluate the sensitivity of novel antibody-conjugated fluorochromes in near-infrared region. Compensation beads were stained with each novel antibody-conjugated fluorochromes and acquired either on the APD-based spectral flow cytometer Cytek Aurora or the PMT-based flow cytometer BD FACSymphony A5. As for BD FACSymphony, 710/50, 780/60, 820/60 bandpass filters on both blue laser and red laser were used for acquisition. Stain index from each measurement of novel antibody-conjugated fluorochromes was calculated as previously described^6^.

### Stability test of the novel fluorochromes after fixation

Peripheral blood mononuclear cells (Veri-Cells PBMC, BioLegend) were stained with each novel fluorochrome for 30 minutes and fixed with various fixatives according to the manufacturer’s instructions – Cell Staining Buffer (PBS + 10% Fetal Bovine Serum (FBS)), FACS lysing solution (BD Biosciences), Fixation buffer and Intracellular Staining Perm Wash buffer (BioLegend), and Foxp3/Transcription factor staining buffer with Permeabilization buffer (Thermo Fisher). Samples were analyzed directly after fixation and washing (marked as “fresh”) and analyzed after overnight incubation at 4°C (marked as “Overnight”). All samples were acquired on Aurora spectral flow cytometry and stain index of each fluorochrome was calculated.

### Whole blood staining of immune cell subsets for flow cytometry

Monocyte blocking solution (BioLegend) and Brilliant Stain Buffer (BD) were added prior to multicolor staining. A cocktail of novel in-house antibody-conjugated fluorochromes and other antibody-conjugated fluorochromes was added to 300 μl of whole blood in a 12 × 75 mm tube. Single-color reference controls (also known as single color compensation bead controls) were stained with each antibody-conjugated fluorochrome. 0.5 - 1 μg of in-house antibody-conjugated fluorochromes per test was used. 5 μl of commercially available antibody-conjugated fluorochromes per test was used as instructed. 1 μl of HLA-DR-APC-eFluor780 and 0.25 μl of CD4-BB660 per test was used due to its intense brightness.

Antibody-conjugated fluorochromes for the violet 405nm laser includes: CD33-BV421 (Clone WM53), CD28-BV510 (Clone 28.2), CD27-BV570 (Clone O323), CD123-BV650 (Clone 7G3), CXCR5-BV711 (Clone J252D4), CD56-BV750 (Clone 5.1H11) (BioLegend); CD22-SuperBright436 (Clone 4KB128), CD57-eFluor450 (Clone TBO1), Qdot 585 Streptavidin conjugate (Invitrogen); CD138-BV480 (Clone MI15), CD127-BV605 (Clone HIL-7R-M21), CD45RA-BV786 (Clone HI100) (BD); CD14-KromeOrange (Clone RMO52) (Beckman Coulter).

Antibody-conjugated fluorochromes for the blue 488nm laser includes: DNAM-BB515 (Clone DX11), CXCR3-PE (Clone 1C6/CXCR3), CCR7-PE-CF594 (Clone 150503), CD4-BB660 (Clone SK3) (BD); IgD-FITC (Clone IA6-2), CD11b-Alexa Fluor 532 (Clone M1/70), CD11c-PE-Cy5 (Clone 3.9), CD38-PerCP-eFluor710 (Clone 90) (Invitrogen); PD1-PC5.5 (Clone PD1.3) (Beckman Coulter); CCR4-PE-Vio770 (Clone REA279) (Miltenyi Biotec, Auburn, CA).

Antibody-conjugated fluorochromes for the red 640 nm laser includes: TCRγδ-APC (Clone B1), CD303-AlexaFluor647 (Clone 201A) (BioLegend); CD25-APC-R700 (Clone 2A3) (BD); HLA-DR-APC-eFluor780 (Clone LN3) (Invitrogen).

The samples were first incubated with CXCR3, CCR7, and CCR4 for 10 minutes at 37 °C and incubated with rest of the antibodies for 30 minutes in the dark at room temperature. 2ml of 1 × FACS lysing solution (BD, San Jose, CA) were added to the mixture and incubated for another 10 minutes in the dark at room temperature. The tubes were centrifuged at 500 × g for 5 minutes. Qdot 585 VIVID streptavidin conjugate (Thermo Fisher, Carlsbad, CA) was added to the samples and then incubated for 15 minutes. The samples were washed with 2 ml of FACS buffer (1 × PBS with 10% FBS) and centrifuged at 500 × g for 5 minutes. Washing step was repeated twice. In the end, the samples were resuspended in 350μl of FACS buffer.

### Flow Cytometry and High-dimensional Data Analysis

The samples and single-color controls were acquired on an Aurora spectral flow cytometer (Cytek Biosciences, Fremont, CA) with custom configuration (Supplementary Table 1) at Abbvie Biotherapeutics (Redwood City, CA). Using single-color controls and voltage titration method, voltage of each channel was adjusted for optimal sensitivity^8^. Sample QC and unmixing was run on SpectroFlo.

Acquired data was analyzed using FlowJo analysis software (BD Biosciences). viSNE plots and FlowSOM plots were created in Cytobank (www.cytobank.org). All pah rameters were displayed with an arcsinh transformation.

## RESULTS

### Conjugation of protein dye with a small molecule dye

Novel tandem fluorochromes were generated by a principle of Fluorescence Resonance Energy Transfer (FRET). Distance-dependent energy transfer from a donor molecule to an acceptor molecule creates a unique emission wavelength. Here we used PE and APC as a donor molecule due to their high solubility, brightness, and stability. Also, a variety of acceptor dyes were evaluated and highly water-soluble fluorescent dyes were selected as acceptor dyes for covalent labeling.

Tandem dyes were synthesized by a reaction between the NHS ester on the small molecule dye and the amine residue on the protein dye. As previously described^5^, for each tandem dye, various molar ratios of a protein dye and a small molecule dye (ranging from 1:10 to 1:80) were evaluated. By comparing relative brightness, residual donor emission, and emission spectrum at distinctive wavelength, optimal ratio for each tandem dye was determined (Supplementary Table 3). Depicted in Table 1 is the optimal reaction condition for each tandem dye (Table 1). Overall, PE tandems tend to require higher molar ratio supposedly due to its higher molecular weight.

The excitation and emission wavelength were measured (Figure 1). When APC and PE were conjugated with Dy704 and Dy704-P4 respectively, their peak emission wavelength was 750 nm. When APC or PE was conjugated with Dy800-P4, the peak emission wavelength was 800 nm. Conjugates prepared with iFluor810 showed a peak emission wavelength of 830 nm in near-infrared range. We named these novel fluorochromes as a combination of donor molecule and its peak emission wavelength: PE-750, PE-800, PE-830 and APC-750, APC-800, APC-830.

**Figure 1.**
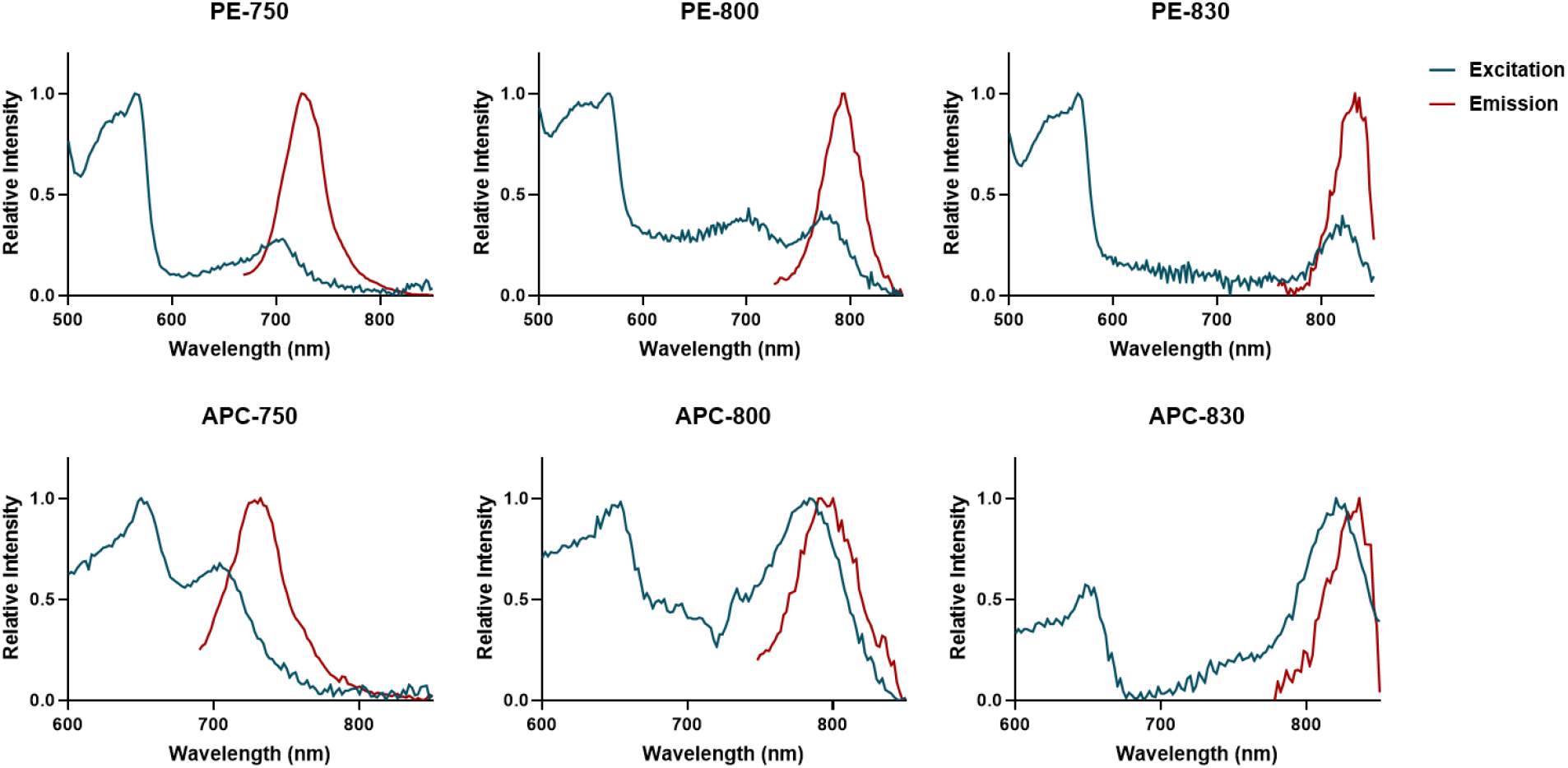
Excitation and emission curves of the novel fluorochromes.

### Conjugation of novel-fluorochromes to a monoclonal antibody

The novel fluorochromes are conjugated with antibodies using click chemistry reaction^7^. Click chemistry is a simple and robust reaction that is commonly used in bioconjugation. It generates conjugated product with quick, high-yield, and high-reaction specificity.

First, an antibody was linked to the TCO tag, and the novel fluorochromes were linked to methyl-tetrazine, respectively. Then, the antibody-TCO structure and fluorochrome-methyl-tetrazine were crosslinked by mixing two reagents in 1:2 molar ratio, respectively. The reaction is completed in 1-2 hours at room temperature or overnight at 4°C.

We selected lineage markers for antibody conjugation with the novel fluorochromes. Their expression pattern is very predictable, so the quality of the novel antibody-conjugated fluorochromes can be easily evaluated.

Initial assessment of the Cytek Aurora reveals the following available channels: B11, B14, B16 for the blue laser and R5, R8, R10 for the red laser (Supplementary Table 2). The fluorochromes that we created demonstrated peak emission spectra at the desired channels on the Cytek Aurora (Figure 2). We further performed stability testing on the final conjugates and found their stability and brightness at 6-month post-conjugation.

**Figure 2.**
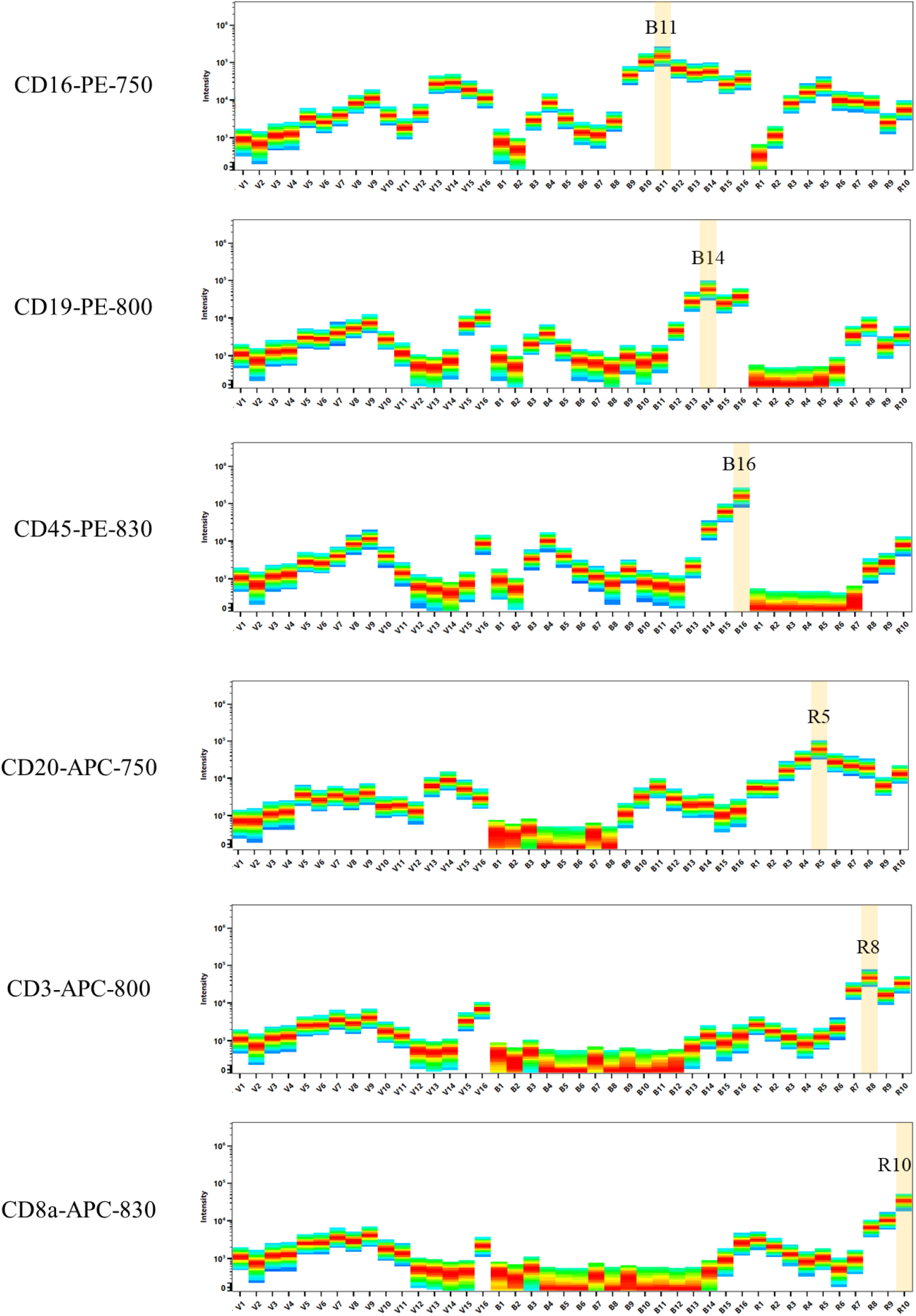
Spectral plots of the novel antibody-conjugated fluorochromes. Each spectral plot shows its peak emission and spectrum signatures throughout violet, blue, and red lasers.

### Comparison of sensitivity between APD-based and PMT-based flow cytometers, and assessment of the novel antibody-conjugated fluorochromes stability in fixatives

PMT-based system has been widely used in flow cytometry. Here, APD technology in spectral flow cytometry was compared against the PMT-based system, specifically for performance in the near-infrared region.

On the FACSymphony system, 710/50 bandpass filters off the blue laser (488 nm) and red laser (640 nm) were used to measure PE-750 and APC-750 fluorescent signals. PE-800 and APC-800 were measured with the 780/60 bandpass filters, while 820/60 bandpass filters were used to measure PE-830 and APC-830 fluorescent signals.

Not surprisingly, the Cytek Aurora demonstrated higher staining index for all fluorochromes, except for APC-750, over the BD FACSymphony (Figure 3A). The differences in the performance were the most striking for PE-830 and APC-830, validating that APD indeed have superior sensitivity over PMT for emission wavelength beyond 800nm.

**Figure 3.**
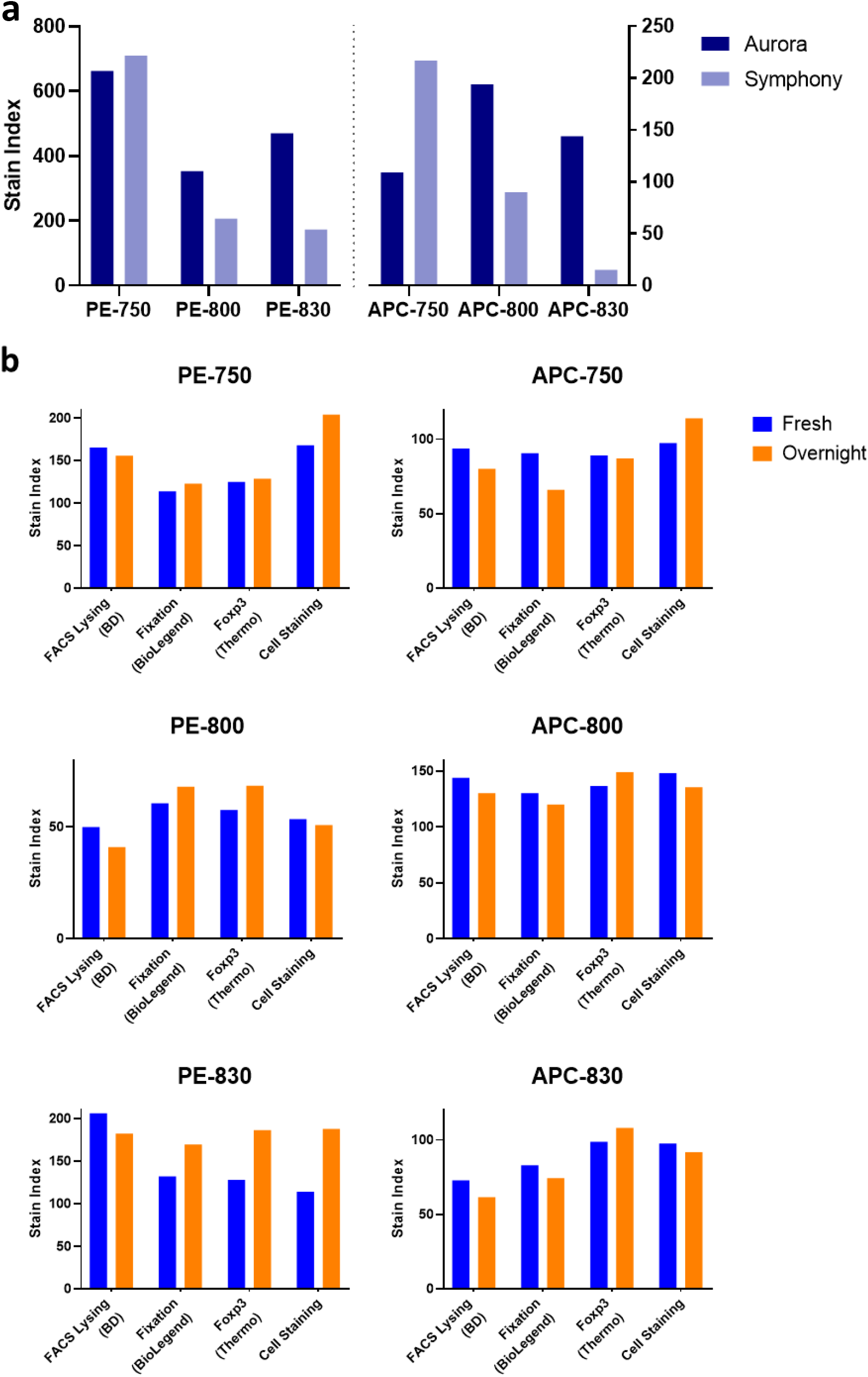
Novel antibody-conjugated fluorochromes showed higher sensitivity in near-IR region in spectral flow cytometry and maintained stability after fixation followed by storage overnight. Stain indexes were obtained for each of the novel antibody-conjugated fluorochromes after binding to compensation beads (Fig 3a) and acquired using Aurora (blue columns) and BD FACSymphony (orange columns). Stability for each antibody-conjugated fluorochrome (Fig 3b) was evaluated in the presence of BD FACS Lysing Solution, BioLegend Fixation Buffer, Foxp3/Transcription factor staining buffer set from Thermo Fisher, and Cell Staining Buffer (PBS + 10% FBS), respectively, by comparing the stain indexes for PBMCs when stained and acquired fresh (blue columns) versus following storage 4°C overnight (orange columns).

Fluorochrome stability after fixative treatment was evaluated in Figure 3B. We selected 3 representative fixative buffer systems that are commonly used in flow cytometry: BD FACS Lysing Solution (1% paraformaldehyde and 3% diethylene glycol, commonly used for whole blood cell surface staining), Biolegend Fixation Buffer (4% paraformaldehyde, commonly used for intracellular cytokine staining), and Thermo Fisher Foxp3 Transcription Factor Buffer kits (proprietary fixative formula, commonly used for transcription factor staining). Single-color cell staining of each novel fluorochrome conjugated with anti-CD4 antibody was either fixed and immediately acquired on the cytometer, or fixed overnight at 4°C and acquired on the next day. As a control, we also stained the cells and resuspend it in cell staining buffer (PBS + 10% FBS). In general, the stain index of the novel antibody-conjugated fluorochromes was not significantly changed after overnight incubation with various fixatives. Also, stain index was similar among treatment with different fixatives. These results suggest that these novel-fluorochromes can be adopted in protocols including fixation and permeabilization steps.

### Spillover spread matrix (SSM) including the novel antibody-conjugated fluorochromes

Spillover spread matrix (SSM) is a useful tool to characterize dye and instrument performance, which helps cytometrist choosing optimal fluorochrome combinations in a panel design. Commonly used 21 fluorochromes conjugated with anti-CD4 antibody in addition to the 6 novel fluorochromes were used to generate the spread matrix in FlowJo, as previously described^8^. SSM value lower than 2 is considered as minimal spread (green), between 2 and 6 as moderate spread (yellow), and above 6 as high spread (red) (Figure 4). As demonstrated by the SSM matrix, PE-800, PE-830, APC-800, APC-830 fluorochromes introduce minimal spread into other detectors, indicating that they could be easily adopted into an existing multicolor panel. In contrast, PE-750 and APC-750 introduce a significant level of spread into other detectors, indicating that thoughtful panel design is needed to incorporate these fluorochromes into an existing multicolor panel.

**Figure 4.**
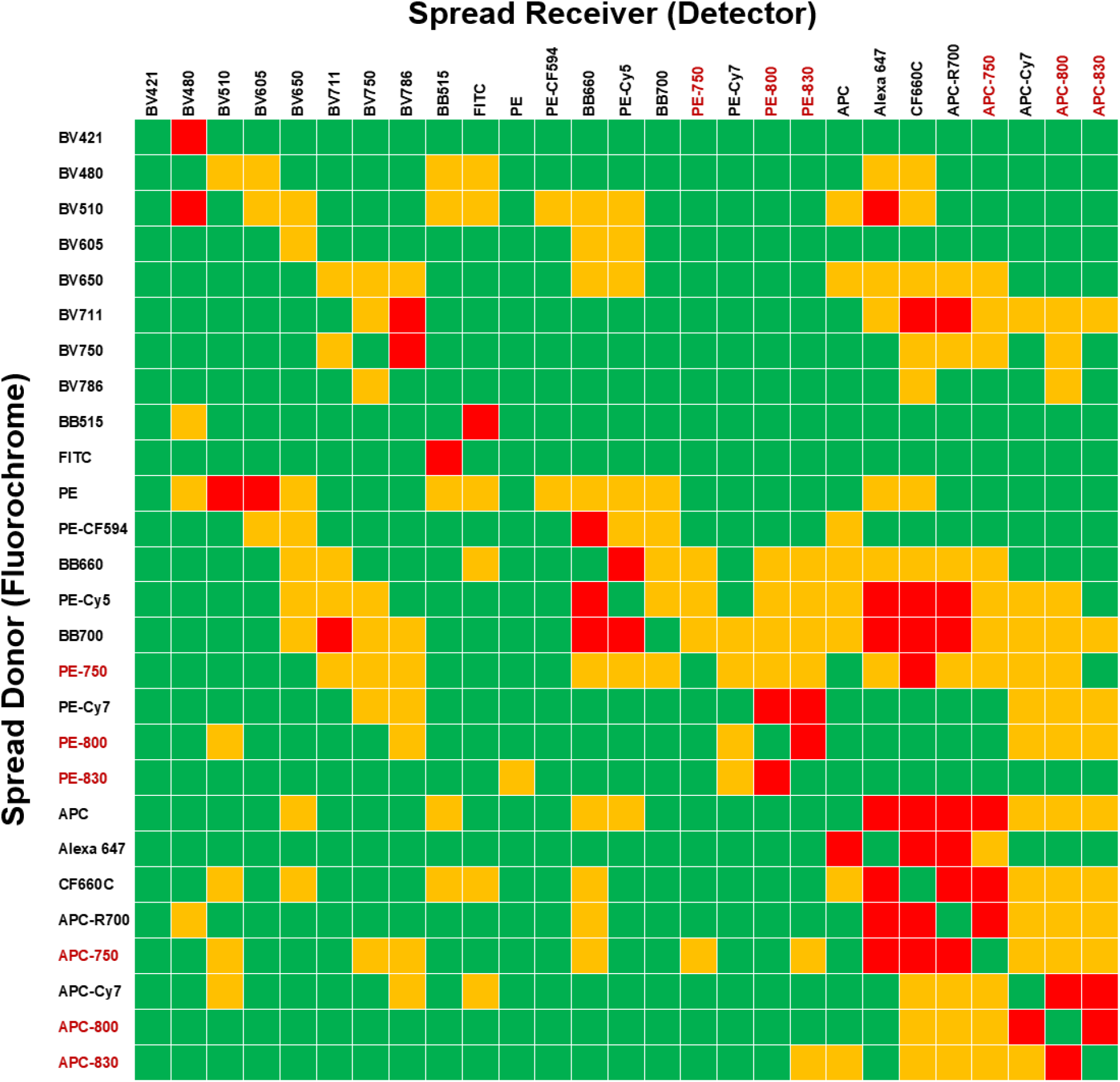
Spillover spread matrix (SSM) calculated with the reagents listed in the first column on spectral flow cytometry. SSM values are color-coded by green-yellow-red color transition; SSM values < 2 (green), between 2 and 6 (yellow), and 6 > (red). The novel antibody-conjugated fluorochromes are written in red.

### An example of a high-dimensional flow cytometry panel incorporating the novel fluorochromes

To demonstrate the utility of the novel fluorochromes, we attempted to design a 34-color panel for broad human immunophenotyping by incorporating 6 novel reagents along 28 commercially available reagents (**Table 2**).

**Table 2.** 34-color flow cytometry panel for broad immunophenotyping in human. 6 novel antibody-conjugated fluorochromes were combined with 28 commercially available antibody reagents – 13 channels for violet laser, 13 channels for blue laser, and 8 channels for red laser. Novel in-house antibody-conjugated fluorochromes are marked in yellow.

| Violet |  | Blue |  | Red |  |
| --- | --- | --- | --- | --- | --- |
| Fluorochrome | Antibody | Fluorochrome | Antibody | Fluorochrome | Antibody |
| BV421 | CD33 | BB515 | DNAM | APC | TCR$\gamma\delta$ |
| SB436 | CD22 | FITC | IgD | Alexa Fluor 647 | CD303 |
| eFluor 450 | CD57 | Alexa Fluor 532 | CD11b | CF680 | CCR6 |
| BV480 | CD138 | PE | CXCR3 | APC-R700 | CD25 |
| BV510 | CD28 | PE-CF594 | CCR7 | APC-750 | CD20 |
| Krome Orange | CD14 | BB660 | CD4 | APC-eFluor780 | HLA-DR |
| BV570 | CD27 | PE-Cy5 | CD11c | APC-800 | CD3 |
| Qdot585 | CCR3 | PE-Cy5.5 | PD1 | APC-830 | CD8 |
| BV605 | CD127 | PerCP-eFluor710 | CD38 |  |  |
| BV650 | CD123 | PE-750 | CD16 |  |  |
| BV711 | CXCR5 | PE-Vio770 | CCR4 |  |  |
| BV750 | CD56 | PE-800 | CD19 |  |  |
| BV786 | CD45RA | PE-830 | CD45 |  |  |

To optimize the instrument settings for the novel fluorochromes, the voltages for each channel were adjusted by the voltage titration method to maximize resolution (Supplementary Table 1) as previously described^9,10^. After setting the voltages, reference controls (also known as single color compensation bead controls) were acquired and used for spectral data unmixing. In some cases, we recognize that compensation beads may behave differently from cells. Therefore, we fine-tuned the compensation matrix by imposing the compensation bead-derived matrix on single-color stained blood cells and made minor adjustment to the matrix to correct for the differences between cells and compensation beads. The final matrix was applied to the samples. Single staining of each novel antibody-conjugated fluorochrome is shown in Supplementary Figure 1. The final unmixing matrix was applied to the blood donor samples.

To demonstrate the general utility of the 34-color flow panel, we simply gated major immune cell subsets using 2D dot plots in SpectroFlo – the default analysis software package on the Aurora cytometer (Figure 5). Within single lymphocytes, central memory CD4 and CD8 T cells were gated as CD4^+^CD27^+^CD45RA^-^ and CD8^+^CD27^+^CD45RA^-^, respectively. Regulatory T cells were gated as CD4^+^CD25^+^CD127^-^ cells. Within CD3^-^ lymphocytes, we identified memory B cells as CD20^+^CD27^+^IgD^-^ and naïve B cells as CD20^+^CD27^-^IgD^+^. Similarly, within the CD3^-^CD20^-^ lymphocytes, NK cells were divided into two groups CD56^+^CD16^-^ and CD56^-^CD16^+^ NK cells. Rare populations such as plasma cells (CD38^++^ and CD138^+^) and plasmacytoid dendritic cells (CD303^+^ and CD123^+^) could be easily identified.

**Figure 5.**
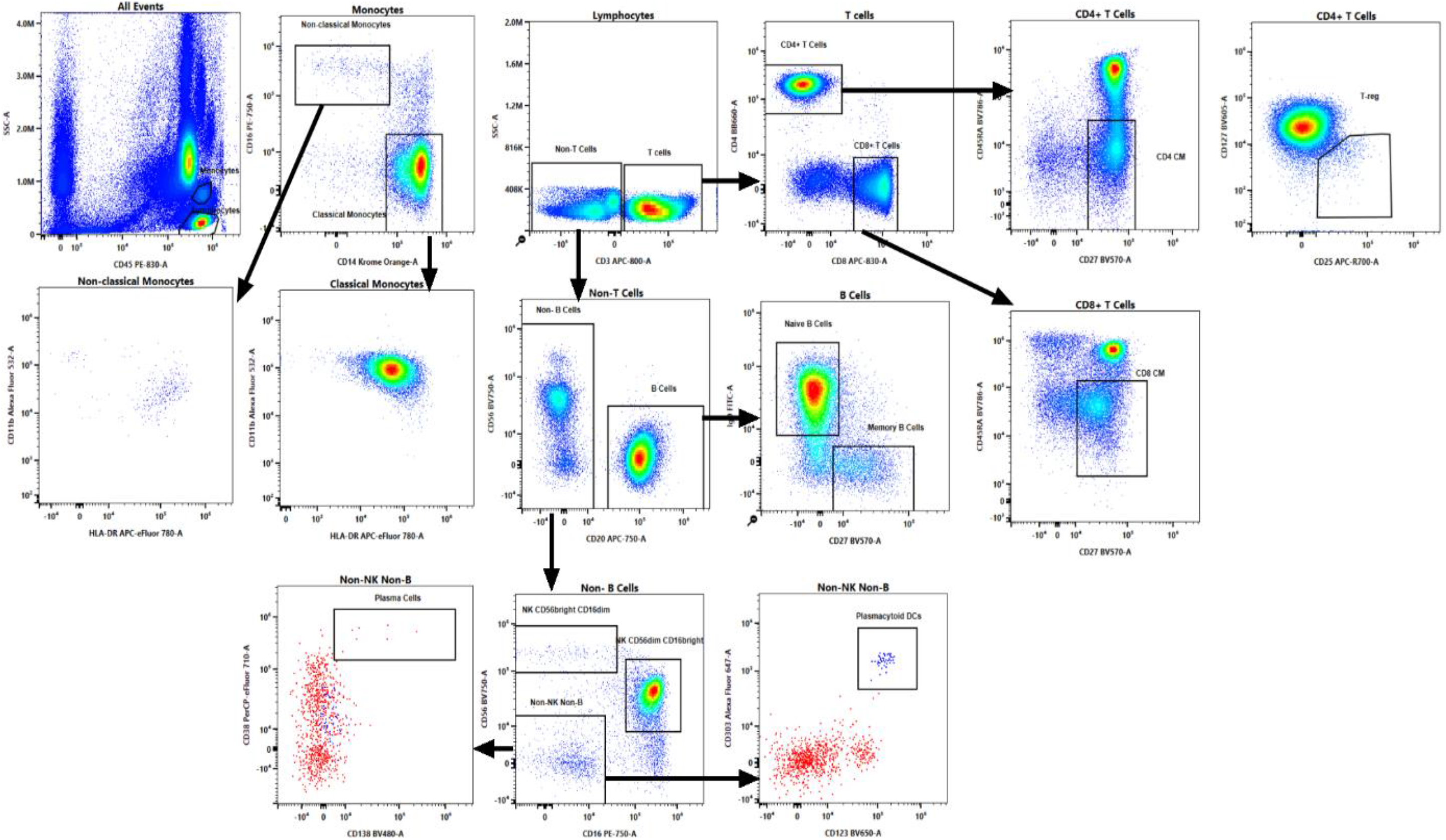
2D dot plots for immune cell subsets in human whole blood. Some immune cell subsets were gated by cell-lineage markers. CM: central memory; Treg: regulatory T cells.

### High dimensional analysis using viSNE confirmed antibody staining patterns on individual cells was consistent with established immune phenotype expression patterns

A variant of t-Distributed Stochastic Neighbor Embedding (t-SNE) analysis, termed viSNE, was performed on results obtained following antibody cocktail staining of blood from three human donors (Supplementary Figure 2) using an antibody cocktail that incorporated one of each of the novel fluorochromes as well as 28 additional cell surface antigen-specific fluorescently labeled antibodies. These viSNE graphic visualizations confirmed that staining of each antibody in the cocktail was specific for immune cells known to express each of those antigens. For example, CD3 staining intensities were highest for CD4^+^ and CD8^+^ cells, but not detectable for CD20^+^ cells and PD-1 levels were highest on immune cells with a CD4 effector memory T cell phenotype (CD4^+^CD45RA^-^CD27^-^) and lower on cells with a CD4 naïve (CD4^+^CD45RA^+^CD27^+^) phenotype. Following viSNE analysis, FlowSOM could be used to generate clusters and provide statistical analysis to identify the most significant population cluster node (Supplementary Figure 3). These analyses may accelerate novel biomarker discovery using high-content flow cytometry panels in clinical and pre-clinical studies.

## DISCUSSION

Recent advances of flow cytometric instrumentation have significantly increased the number of parameters that can be measured simultaneously at the single cell level. Here we aimed to expand the capacity of a common flow cytometer platform by creating fluorochromes for channels where commercially available antibody conjugates are limited. Six combinations of tandem fluorochromes were generated and were successfully conjugated to monoclonal antibodies for characterization. Antibody conjugation to each novel fluorochrome was achieved using simple Click Chemistry reactions.

The SSM shows that these novel antibody-conjugated fluorochromes can be easily combined with other commercially available fluorochromes. Stability test with fixative buffers showed that they can be used for experiments requiring fixation and permeabilization steps.

Recently various groups have demonstrated examples of 40-color flow cytometry panel run on the 5-laser Aurora^11,12^. In this study, we designed and achieved a 34-color panel using Aurora system possessing only 3 lasers by utilizing six novel fluorochromes.

Taken together, these results demonstrate novel fluorochrome development to enable the high sensitivity, simultaneous measurement of high-content flow cytometry. The tandem-dye approach, as demonstrated here, has the potential to provide novel fluorochromes that could expand the number of simultaneous parameters.

## ACKNOWLEDGEMENTS

We thank AbbVie Biotherapeutics Employee Blood Collection Program for providing human blood samples. We would also like to thank James Sheridan of AbbVie for the careful review and feedbacks.

