## Supplemental Figures and Tables for "Novel PE and APC Tandems: Additional Near-Infrared Fluorochromes for Use in Spectral Flow Cytometry"

### SUPPLEMENTARY FIGURES AND TABLES

| | Violet | Blue | Red |
|---|---|---|---|
| 1 | 511 | 1492 | 450 |
| 2 | 729 | 397 | 525 |
| 3 | 804 | 318 | 821 |
| 4 | 512 | 215 | 801 |
| 5 | 3069 | 196 | 932 |
| 6 | 955 | 344 | 533 |
| 7 | 846 | 589 | 559 |
| 8 | 848 | 519 | 908 |
| 9 | 1050 | 653 | 292 |
| 10 | 974 | 777 | 518 |
| 11 | 589 | 794 | |
| 12 | 437 | 649 | |
| 13 | 410 | 567 | |
| 14 | 547 | 792 | |
| 15 | 686 | 294 | |
| 16 | 534 | 817 | |

**Supplementary Table 1. Voltage setting for each channel of 3-laser Aurora system.**

| Laser | Channel | Center Wavelength (nm) | Bandwidth (nm) | Wavelength Start (nm) | Wavelength End (nm) | Corresponding Fluorochromes |
|---|---|---|---|---|---|---|
| Blue | B1 | 508 | 20 | 498 | 518 | BB515 |
| | B2 | 528 | 21 | 518 | 539 | FITC / AF488 |
| | B3 | 549 | 22 | 538 | 560 | AF 532 |
| | B4 | 571 | 23 | 560 | 583 | PE |
| | B5 | 594 | 23 | 583 | 606 | |
| | B6 | 618 | 24 | 606 | 630 | PE-CF594 |
| | B7 | 660 | 17 | 652 | 669 | PE-Cy5 |
| | B8 | 678 | 18 | 669 | 687 | PerCP-Cy5.5 |
| | B9 | 697 | 19 | 688 | 707 | PerCP-eFluor710 |
| | B10 | 717 | 20 | 707 | 727 | |
| | B11 | 738 | 21 | 728 | 749 | |
| | B12 | 760 | 23 | 749 | 772 | |
| | B13 | 783 | 23 | 772 | 795 | PE-Cy7/PE-Vio770 |
| | B14 | 812 | 34 | 795 | 829 | |
| | B15 | 843 | 30 | 829 | 858 | |
| | B16 | 871 | 25 | 859 | 884 | |

| Laser | Channel | Center Wavelength (nm) | Bandwidth (nm) | Wavelength Start (nm) | Wavelength End (nm) | Corresponding Fluorochromes |
|---|---|---|---|---|---|---|
| Red | R1 | 660 | 17 | 652 | 669 | APC |
| | R2 | 678 | 18 | 669 | 687 | AF647 |
| | R3 | 697 | 19 | 688 | 707 | CF680 |
| | R4 | 717 | 20 | 707 | 727 | APC-R700 |
| | R5 | 738 | 21 | 728 | 749 | |
| | R6 | 760 | 23 | 749 | 772 | |
| | R7 | 783 | 23 | 772 | 795 | APC-Cy7/APC-eFlour780 |
| | R8 | 812 | 34 | 795 | 829 | |
| | R9 | 843 | 30 | 829 | 858 | |
| | R10 | 871 | 25 | 859 | 884 | |

**Supplementary Table 2. Wavelength range properties of each channel of blue and red lasers.** Compared to a standard Aurora, the special order Aurora spectral flow cytometer in AbbVie, Redwood City has additional channels (B15, B16, R9, and R10) for blue and red lasers to detect peak emission in the near-infrared regions. Channels where the novel fluorochromes fit colored blue or red.

| APC-800 (APC-Dy800P4) Reaction Input | Channel 1 MFI (R1) | Channel 2 MFI (R8) | Brightness Ratio (R8/R1) | Compensation % |
|---|---|---|---|---|
| **1:10** | **859** | **23988** | **27.93** | **3.581** |
| **1:15** | **356** | **21409** | **60.14** | **1.663** |
| **1:20** | **77.7** | **8151** | **104.90** | **0.953** |

| APC-830 (APC-iFluor810) Reaction Input | Channel 1 MFI (R1) | Channel 2 MFI (R10) | Brightness Ratio (R10/R1) | Compensation % |
|---|---|---|---|---|
| **1:20** | **1758** | **6734** | **3.83** | **26.106** |
| **1:25** | **977** | **5675** | **5.81** | **17.216** |
| **1:30** | **699** | **4892** | **7.00** | **14.289** |

**Supplementary Table 3. Optimal ratio between donor and acceptor dyes were holistically determined based on peak emission MFI, the brightness ratio, and compensation percentages.** Here we show an example of the selection process for an optimal F:P donor:acceptor ratio. Tandem dyes were created by conjugating donor dyes and acceptor dyes at various ratio ranging from 1:10 to 1:80. At the end of the reaction, we measured the peak emission MFI (channel R8 for APC-800 and channel R10 for APC-830) and the residual donor emission MFI (channel R1 for APC-based tandem dyes). We further calculated brightness ratio and determined compensation percentages. Based on these parameters, we holistically decided on the optimum reaction ratio for each tandem dye.

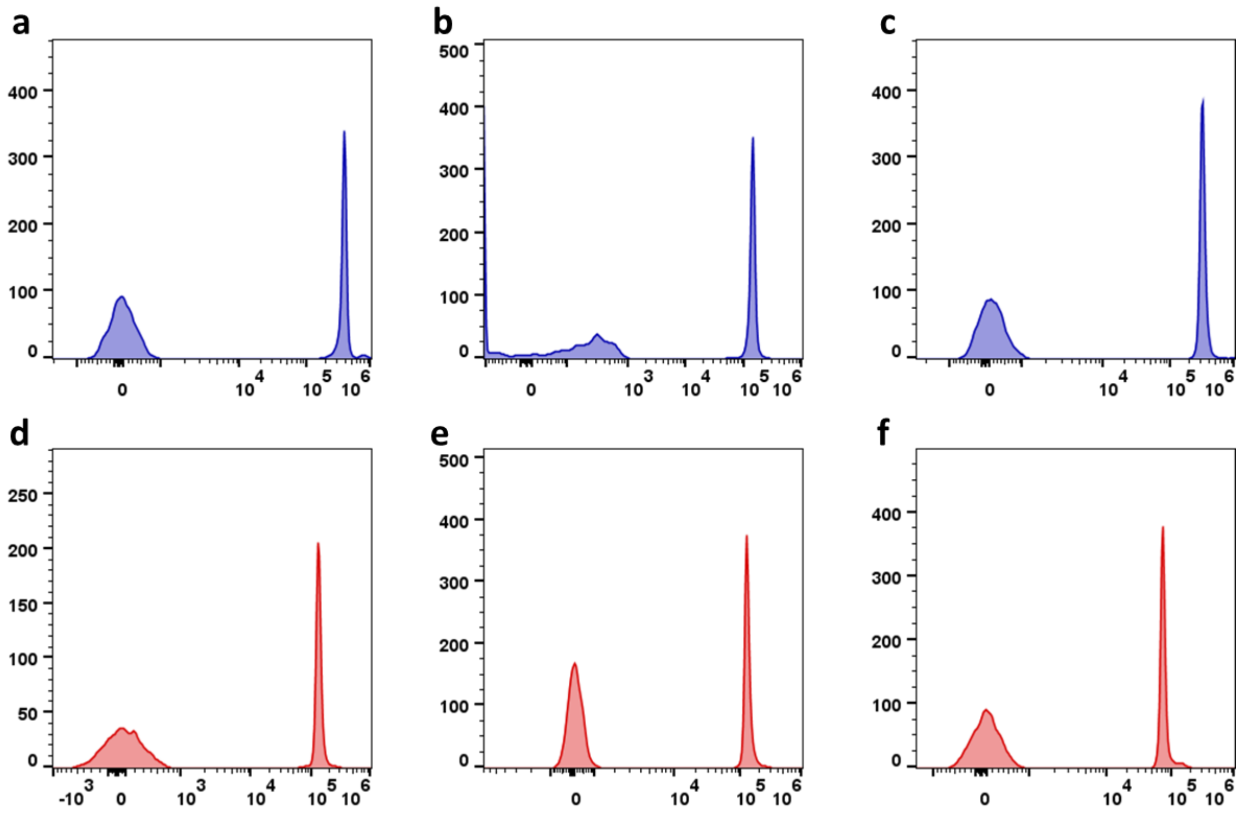


**Supplementary Figure 1. Single staining of the novel antibody-conjugated fluorochromes.** Compensation beads were stained with each antibody-conjugated fluorochrome and acquired by the spectral flow cytometer. The histograms were generated on FlowJo. a. PE-750; b. PE-800; c. PE-830; d. APC-750; e. APC-800; f. APC-830.

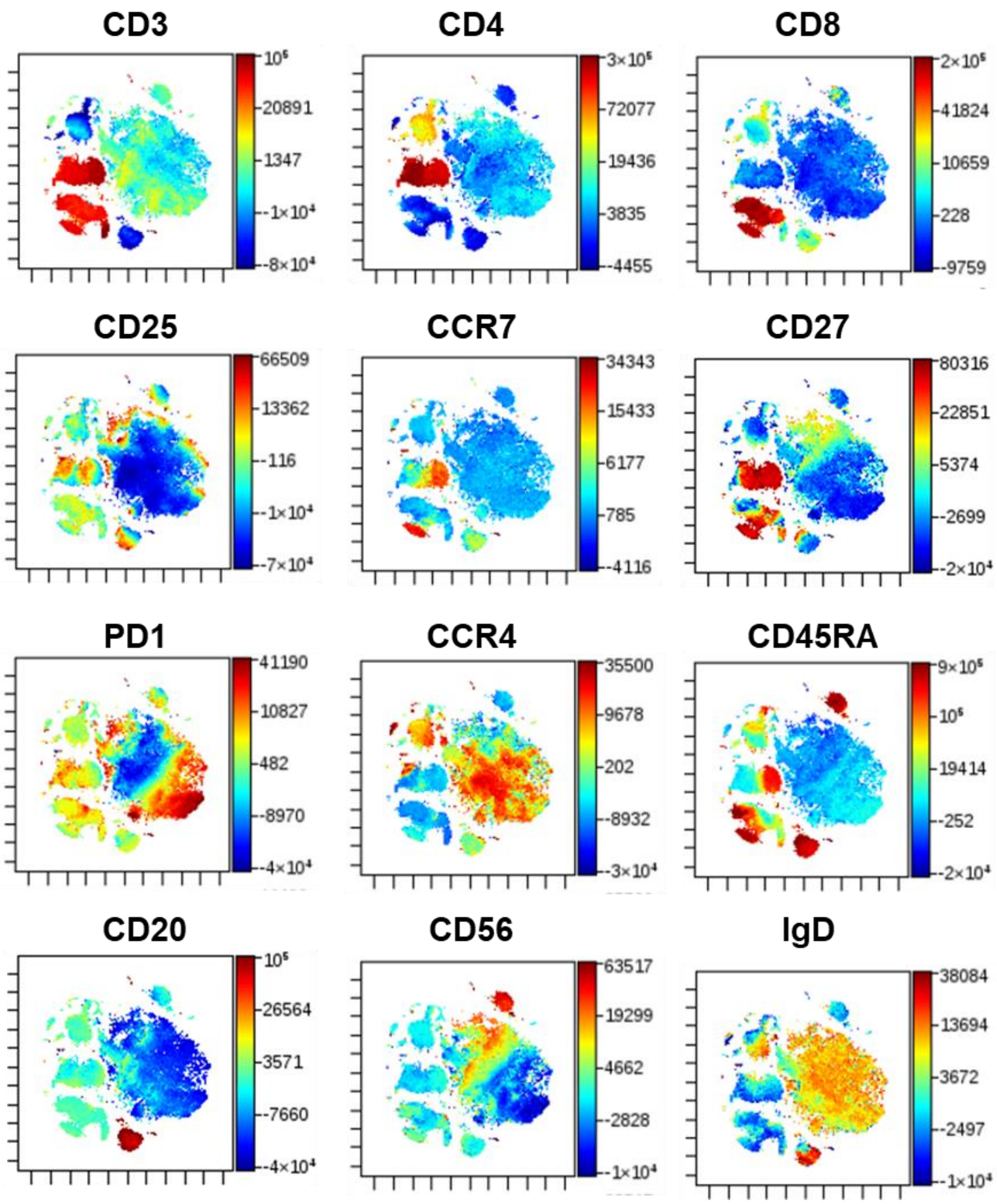


**Supplementary Figure 2. Dimension reduction algorithms visualize expression of each cell type-specific marker.** Reduction of high dimensional datasets into single marker analysis using viSNE confirmed the binding for each antigen specific marker was specific (*e.g., CD3 is highest on $CD4^+$ and $CD8^+$, but not present on $CD20^+$ or $CD56^+$)*. Within live singlets, lymphocytes, monocytes, and granulocytes were used to generate clusters. Perplexity of 70 was used to generate viSNE plots using Cytobank (n=3 donor blood samples).

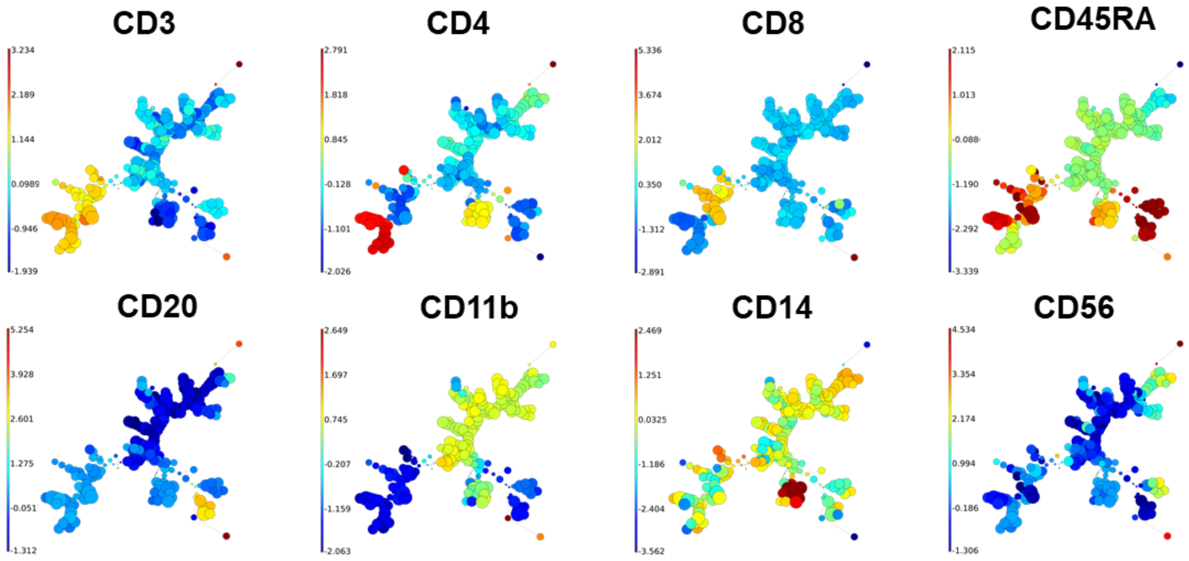


**Supplementary Figure 3. Dimension reduction algorithm FlowSOM generated 15 meta clusters using 3 donor blood samples.** FlowSOM provides a quick and consistent clustering for high-dimensional data by using all markers on all cells using self-organizing map (SOM). Analysis was performed on Cytobank. Some representative plots for lineage markers were shown.
